# Gastrointestinal delivery of mRNA lipid nanoparticles selectively targets the pancreas

**DOI:** 10.1101/2025.09.15.676190

**Authors:** Abdulraouf M. Abbas, Ramy Ghanim, Sebastian Rudden, David Schultz, Avraham Shakked, Hyejin Kim, James E. Dahlman, Alex Abramson

**Affiliations:** Wallace H. Coulter Department of Biomedical Engineering, Emory University School of Medicine and Georgia Institute of Technology, Atlanta, GA, 30332, USA; School of Chemical and Biomolecular Engineering, Georgia Institute of Technology, Atlanta, GA, 30332, USA; Wearable Intelligent Systems and Healthcare Center, Institute for Matters and Systems, Georgia Institute of Technology, Atlanta, GA, 30332, USA; Division of Digestive Diseases, Emory University School of Medicine, Atlanta, GA, 30332, USA

## Abstract

Lipid nanoparticles (LNPs) administered parenterally often show poor localization to the gastrointestinal (GI) tract and pancreas. In addition, patients typically prefer orally administered drugs to those given intravenously. We therefore investigated whether GI delivery, achievable via device mediated microneedle injections applied to buccal, gastric, small intestinal, colonic, or rectal tissues, could simultaneously enhance LNP delivery to the GI and pancreas while avoiding intravenous administration. Using a combined approach of formulation optimization and GI delivery site screening, we found that cationic SM-102 LNPs delivered gastrically achieved 7-fold higher pancreas delivery in rodents than intravenous neutral SM-102 LNPs. With dose optimization, gastric LNPs achieved 6000-fold greater pancreas to liver targeting ratios than intravenous LNPs. These results suggest GI microneedle administration can reprogram LNP biodistribution, thereby expanding therapeutic opportunities for both local and systemic nucleic acid delivery.

## 1. Introduction

Therapeutic nucleic acids feature in over 20 FDA-approved products^1–3^. Messenger RNA (mRNA) drug products have been limited to intramuscularly administered vaccines, while short interfering RNA (siRNA) products have been primarily limited to intravenously and subcutaneously delivered therapies for liver diseases. Non-liver delivery remains challenging, yet nucleic acids show promise in treating preclinical models of gastrointestinal disease including gastroesophageal cancer, gastric carcinoma, and ulcerative colitis as well as pancreatic diseases such as cancer and diabetes ^4–10^. To improve non-liver delivery, labs either pre-treat mice with molecules that inhibit liver uptake^11^ or alter LNP chemistry by adding active targeting ligands or permanently charged helper lipids^12–15^. An alternative strategy involves altering the administration route. Scientists have reported improved lung delivery using nebulization^16^, improved lymphatic delivery using subcutaneous or intramuscular administration^17,18^, and improved pancreas delivery via intraperitoneal administration^19^ yet it remains unclear whether other delivery routes alter LNP tropism.

Given the advent of new technologies for the GI delivery of biologics^20–24^ and evidence that patients prefer oral alternatives to parenteral administration^25,26^, we found this question compelling. Scientists have reported multiple approaches to avoid the harsh environment and diffusional barriers of the GI tract^27^, including endoscopic injections^23^, miniaturized ingestible microneedle pills^21^ and patches^28^, jet injection pills^4,24^, ultrasound mediated enemas^29^, and novel formulations^30–32^. All of these have generated data consistent with local endoluminal and buccal nucleic acid uptake. However, it remains unclear how altering the GI-related route of administration affects delivery systemically; the on- and off-target delivery ratios as a function of administration route are especially important given that unwanted liver delivery is an established challenge in the field of nucleic acid therapeutics.

Here we systematically investigated LNP-mRNA delivery to the submucosa of clinically relevant GI organs including buccal, gastric, small intestine, colonic, and rectal tissue. We discovered that gastric and rectal delivery resulted in 10-fold and 3-fold higher uptake in the pancreas compared to intravenous administration, respectively. This on-target improvement was coupled with significantly lower uptake in off-target organs such as the liver, lungs, heart, and spleen, compared to intravenous administration. Separately, we found that adding a cationic helper lipid to an LNP improved pancreas targeting compared to an LNP formulated with a neutral helper lipid after an intravenous administration. These data demonstrated synergistic effects; the optimized route of administration and LNP composition improved pancreas uptake 7-fold and decreased liver delivery 2.4-fold relative to control LNP delivered intravenously. After further optimizing the dose of a cationic LNP, gastric LNPs achieved 6000-fold greater pancreas to liver targeting ratios than intravenous LNPs. This study highlights that optimizing GI delivery routes and LNP composition may lead to differentiated LNP tropism *in vivo*.

## 2. Results and Discussion

### 2.1 Gastrointestinal LNP administration improves mRNA delivery in the pancreas and decreases mRNA delivery to the liver, lungs, and spleen

We first measured LNP trafficking and delivery as a function of GI-related route of administration. We formulated an LNP using the clinically validated ionizable lipid SM-102 (cite) and the zwitterionic helper lipid 1,2-distearoyl-sn-glycero-3-phosphocholine (DSPC) (termed LNP^neut^) with mRNA encoding anchored nanoluciferase (aNLuc)^33^, then injected them into C57BL/6 mice via the buccal, gastric, colonic, and rectal submucosal tissue in C57BL/6 mice at a dose of 1 mg kg^-1^ (**Figure 1a**). As a control, we administered the LNPs intravenously into mice. Separately, we injected these LNPs into the small intestine of Sprague Dawley rats, since small intestine injections in mice were not feasible (**Figure S1**). LNP^neut^ had a hydrodynamic diameter equal to 78 nm, a 98% encapsulation efficiency, and a polydispersity index (PDI) < 0.3 (**Figure 1b**).

**Figure 1.**
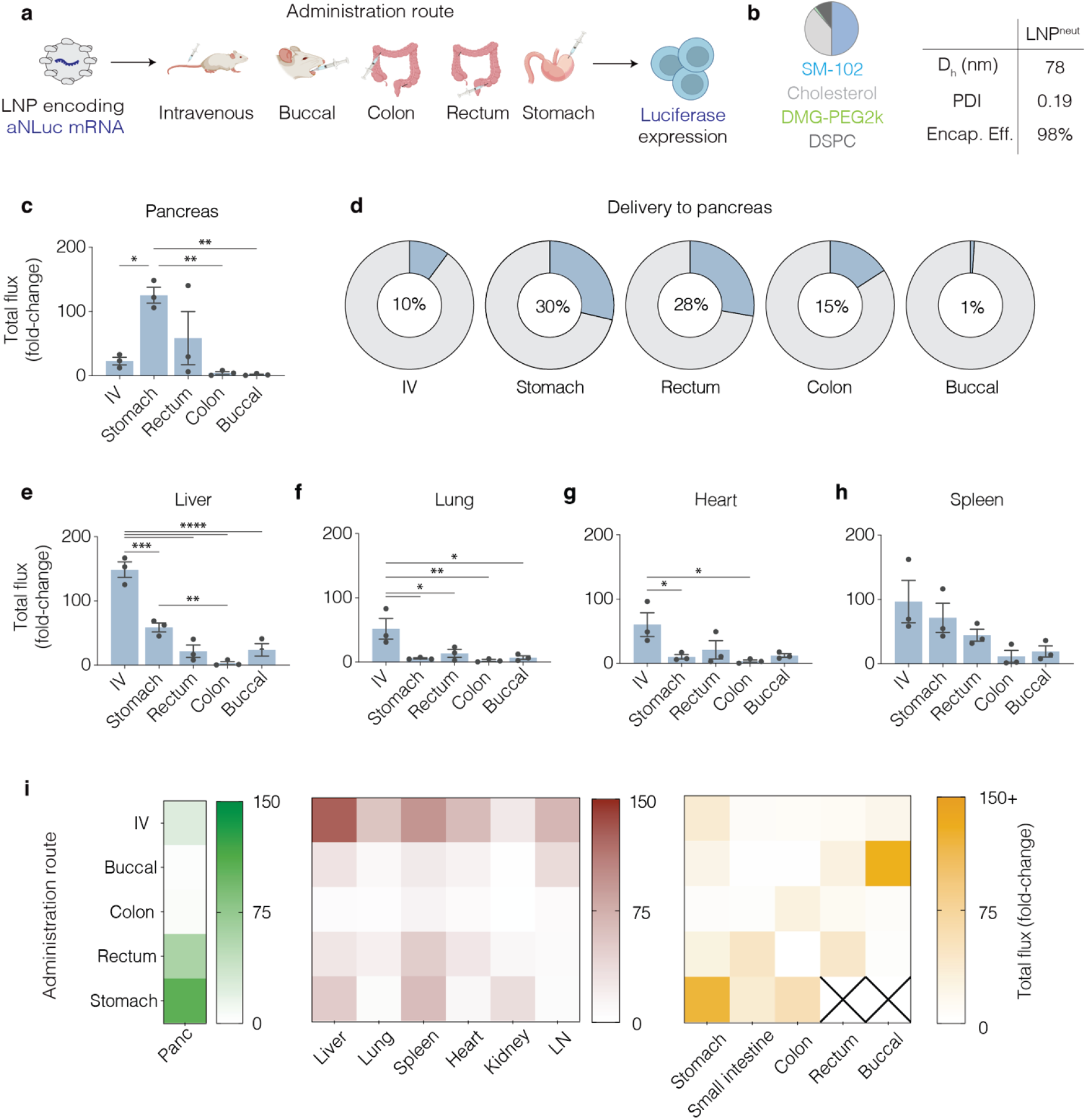
In vivo gastrointestinal delivery of aNLuc mRNA with LNP^neut^ revealed pancreas-targeting upon stomach injection. a) aNLuc RNA was administered at 1 mg kg^-1^ intravenously or into the buccal, colon, rectum, or stomach tissue. IVIS imaging at 18 h following dosing revealed luciferase expression. b) Formulation details of LNP^neut^ and physicochemical characteristics: hydrodynamic diameter (D_h_), polydispersity index (PDI), and encapsulation efficiency. c) Total flux fold-change of pancreas compared to PBS-treated group. d) Percentage of pancreas total flux expression out of 10-organs:(e) liver, (f) lung, (g) heart, and (h) spleen, Stomach (Fig S3a), Colon (Fig S3b), Small intestine (Figure S3c), lymph nodes (Figure S3f), and kidney (Figure S3g). i) Heatmap showing total flux (fold-change) across administration routes of pancreas (green), off-target organs (red), and GI organs (orange). (n=3 mice. Individual data points, Mean +/-SEM. One-Way ANOVA with Tukey’s correction. *p< 0.05; **p<0.01; ***p<0.001; ****p<0.0001).

Eighteen hours after injection we collected the pancreas, spleen, kidney, liver, lung, heart, lymph node, buccal, small intestine, colon, stomach, and rectum. We then quantified luciferase activity via in vivo imaging systems (IVIS) and calculated total flux fold-change for each organ by dividing the total flux (p s^-1^) of an organ from an LNP treated animal by the average PBS-treated counterpart. The stomach injections resulted in the greatest pancreas absolute luminescence fold-change compared to other routes of administration (**Figure 1c**) and the highest percentage of delivery to the pancreas (**Figure 1d**). Also of note, GI organ administration resulted in local uptake at the site of injection, delivering to GI organs not traditionally targeted during intravenous administration (**Figure S2, S3**). By contrast, liver, lung, heart, and spleen luminescence was highest in the intravenous group (**Figure 1e-i**). Similar findings of preferential pancreas targeting have also been reported during intraperitoneal injections. To control for the possibility that LNPs delivered to stomach tissue were leaking out of the stomach into the intraperitoneal space, we injected methylene blue to the stomach, and palpated 50 times with a cotton gauze; we observed no leakage (**Figure S4**). Taken together, we observed that LNP^neut^ delivered to the stomach resulted in 10-fold higher signal in the pancreas compared to IV in addition to a higher pancreas to liver expression ratio. We therefore selected stomach injections as the main GI route of administration for further studies.

### 2.2 LNP chemistry and GI route of administration synergistically improve pancreas delivery

We then evaluated whether formulating an LNP with a cationic helper lipid improved pancreas targeting (**Figure 2a**). To control for the behavior of other LNP components, we simply replaced the zwitterionic DSPC with cationic dimethyldioctadecylammonium (DDAB); we named this LNP^cat^. Similar to LNP^neut^, LNP^cat^ formed LNPs with a 96 nm hydrodynamic diameter, 98% encapsulation efficiency, and a PDI < 0.3 (**Figure 2b**). We then injected C57BL/6 mice with LNP^neut^ and LNP^cat^ carrying mRNA encoding aNLuc at a dose of 1 mg kg^-1^, either via the stomach or intravenously. Eighteen hours later we isolated the same ten organs and analyzed luminescence using IVIS. Both LNP^neut^ and LNP^cat^ when delivered via the stomach compared to intravenous administration (**Figure 2c**). As a control, we compared lung delivery after intravenous administration, and found LNP^cat^ led to more lung delivery than LNP^neut^, as expected based on previous literature^14,15^ (**Figure 2d, S5, S6**). Interestingly, we found that injecting the LNPs into the stomach reduced delivery to the primary targeted organ observed during IV injection and increased delivery to the pancreas. Gastric delivery of LNP^cat^ resulted in 27% greater total-fold flux change than LNP^neut^ delivered intragastrically, and 366% greater total-fold flux change than LNP^neut^ delivered intravenously. When we calculated the ratio of pancreas to liver, lung, heart, and spleen delivery, we found superior pancreas targeting when performing intragastric over intravenous delivery across both LNPs, with intragastrically delivered LNP^cat^ providing the greatest total delivery to the pancreas. (**Figure 2e-h**). These data led us to select LNP^cat^ administered via intragastric injection for further analysis of pancreatic delivery.

**Figure 2.**
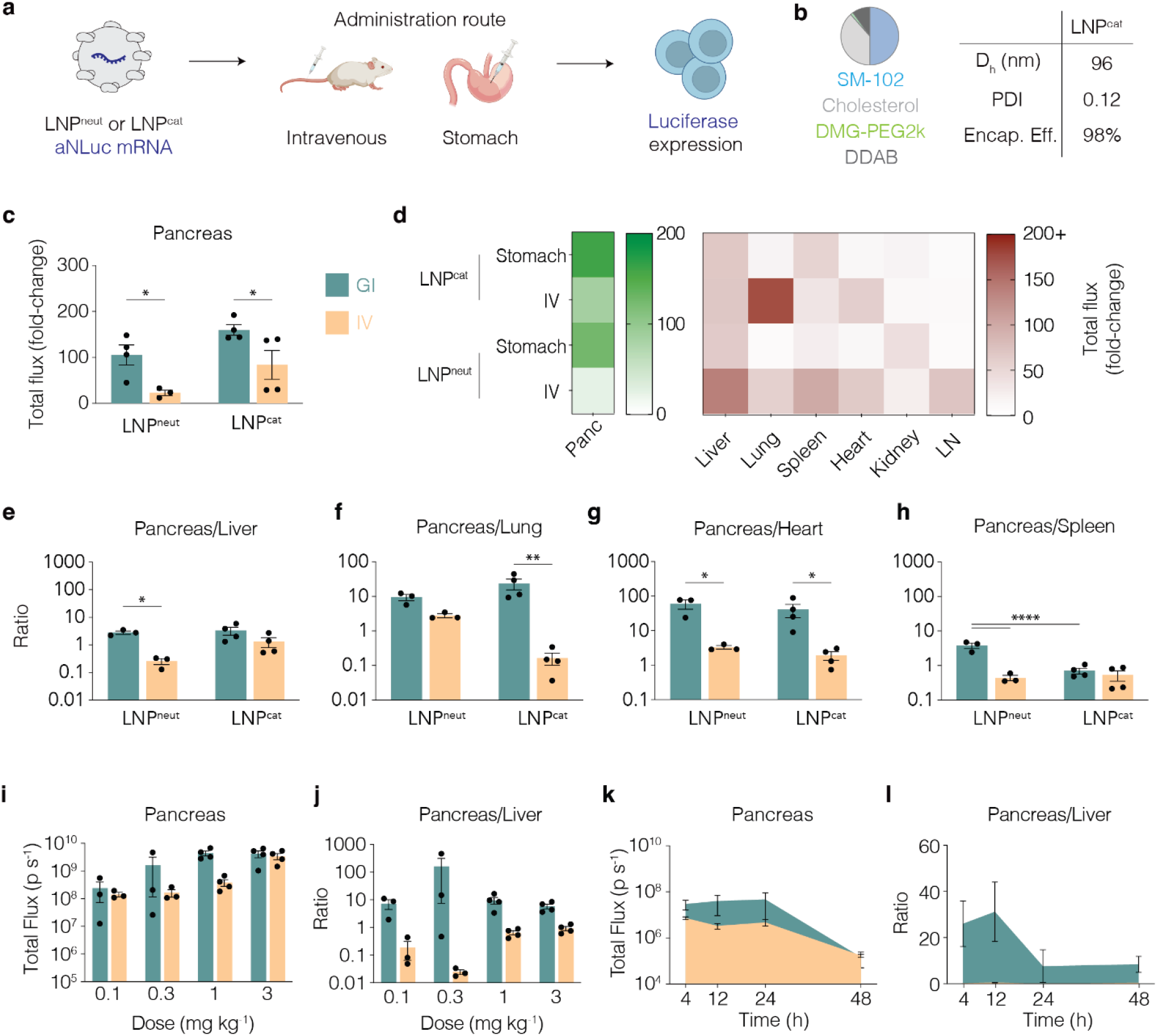
LNP^cat^ increases pancreas-targeting over LNP^neut^, with synergistic targeting effects seen when delivering via stomach compared to IV. a) aNLuc mRNA encapsulated in LNPcat was delivered via IV or stomach at 1 mg kg^-1^. IVIS imaging 18h following dosing revealed luciferase expression. b) Formulation details of LNP^cat^ and physicochemical characteristics: hydrodynamic diameter (D_h_), polydispersity index (PDI), and encapsulation efficiency. c) Total flux (fold-change) of pancreas across IV and stomach administered mRNA with LNP^neut^ vs LNP^cat^ compared to 1x PBS group. d) Heat map showing total flux (fold change) between LNP^neut^ and LNP^cat^ across both IV and stomach administration routes to the pancreas (green), and off-target organs (e) liver, (f) lung, (g) heart, and (h) spleen (red). Additional off target organs in Figure S4. i-j) dose response of LNP^cat^ formulated with aNLuc mRNA 18h post injection showing (i) total flux quantification of pancreas and (j) pancreas/liver ratio. k-l) Time-course of LNP^cat^ formulated with aNLuc mRNA delivered 0.3 mg kg^-1^, at 4h, 12, 24h, 48h. Quantified are (k) total flux of pancreas and (l) pancreas/liver ratios. (n = 3-4 mice. Individual data points, Mean +/-SEM. Two-Way ANOVA with Tukey’s correction. *p<0.05; **p<0.01; ****p<0.0001.)

### 2.3 Gastrointestinal dose response, time course, and safety studies with LNP^cat^

To investigate whether pancreas targeting was observed across a range of doses, we performed a dose-response study. LNP^cat^ was formulated with aNLuc mRNA and administered via IV and gastric injection at 0.1, 0.3, 1.0, or 3.0 mg kg^-1^ in C57BL/6 mice. Eighteen hours after administration, we isolated the pancreas and liver and quantified delivery using IVIS. Pancreas delivery increased with dose, and as expected, saturated at a lower dose after gastric delivery relative to intravenous delivery (**Figure 2i**). Aiming for a maximal pancreas: liver ratio (**Figure 2j**), we chose the 0.3 mg kg^-1^ dose for the subsequent time course study. Specifically, we sacrificed mice 4, 12, 24, and 48 hours after LNP^cat^ administration. Total flux (p s^-1^) in the pancreas was stable for 24 hours, then decreased at the 48-hour timepoint, which is consistent with previous data^34^. As expected from the previous findings, pancreas to liver ratio was minimal for IV throughout this time course, suggesting that therapeutically relevant pancreas delivery can be achieved at lower doses and for longer timescales when delivered intragastrically compared to intravenously. (**Figure 2k, l**). We then assessed safety of LNP^cat^ after a 3 mg kg^-1^ aNLuc mRNA dose administered via gastric injections and intravenously. Specifically, 4 and 20 hours after administration we collected serum, then quantified cytokines and liver damage enzymes. Serum IFNg, TNFa, IL-6, and IL-12p(40) levels from stomach-injected mice were within normal range, and below the IV mouse, at both 4h and 20h timepoints (**Figure S7a-d**). Notably, IFNg and IL-12p40 were elevated in IV mice 4h after administration, but resolved by 20h. Liver enzymes akaline phosphatase (ALP), aspartate aminotransferase (AST), alanine aminotransferase (ALT), and blood urea nitrogen (BUN) were all within normal levels in the stomach injected groups, and all had lower values than IV-injected groups. Only BUN was significantly greater than PBS-injected mice, but still within range (**Figure S7e-h**). These results provide preliminary evidence that administering LNP^cat^ to the stomach is tolerated after a single 3 mg kg^-1^ dose.

### 2.4 Cellular resolution of LNP^cat^ delivery

One limitation to datasets generated with luciferase is the difficulty of determining which cell types are transfected. We therefore performed immunofluorescence (IF) to determine cell types targeted following GI injections. LNP^cat^ was formulated with mRNA encoding Cre recombinase and administered to Ai14 mice via stomach injections at a dose of 1 mg kg^-1^ (**Figure 3a**). Five days after administration we collected pancreatic tissue for histology, and stained with an anti-tdTomato marker as well as hematoxylin and eosin (**Figure 3b**). These data suggested that Cre-mediated genomic excision had occurred in pancreatic acinar cells, pancreatic endothelial cells, and cells within the interlobular duct and the fibrovascular capsule.

**Figure 3.**
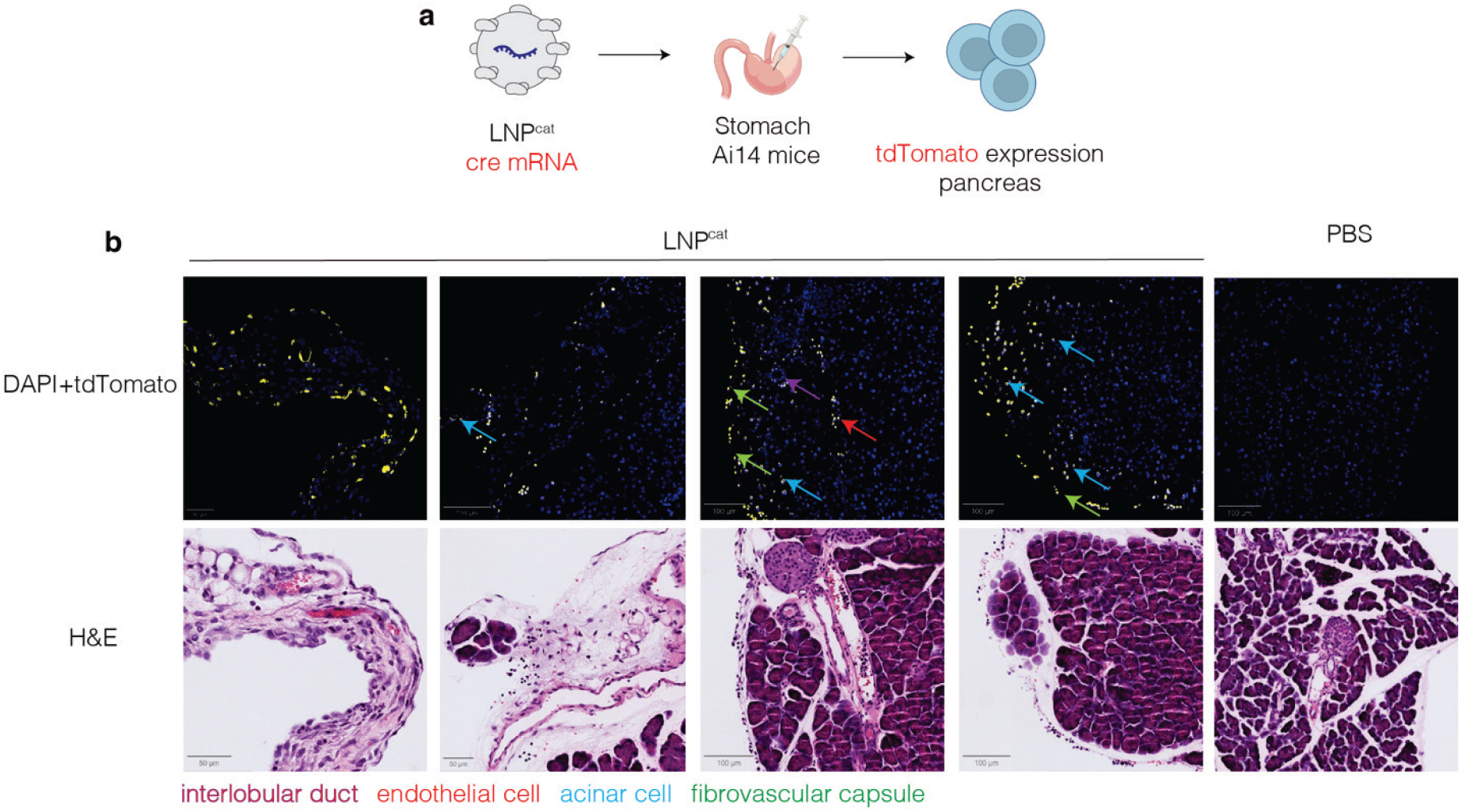
Immunofluorescence targeting tdTomato following delivery of cre mRNA with LNP^cat^ in Ai14 mice shows targeted cell types in pancreas 5 days after delivery. IF and H&E staining shows targeting to multiple cell types.

## 3. Conclusion

In this study, we systematically characterized the biodistributions of LNPs administered via five gastrointestinal delivery routes, including through the submucosal buccal, gastric, small intestine, colonic, and rectal tissues. GI dosing strategies resulted in local uptake that exceeded IV delivery, and gastric administration resulted in pancreas-specific delivery, achieving a 7-fold increase relative to IV dosing. This biodistribution mapping provides evidence of GI dosing as a potential method to reach organs that cannot be targeted parenterally while downregulating hepatic, lung, heart, and splenic expression. GI dosing is especially relevant due to patient preference for oral delivery as well as the recently developed minimally invasive preclinical and clinical robotic capsules being evaluated for delivery of macromolecule formulations to buccal, gastric, and intestinal tissues. Our findings suggest that drug-device combinations could be designed to synergize vehicle formulation strategies with mechanical tissue localization. To fully realize this potential, additional studies in small and large animals are needed to define the interplay between route of administration, particle chemistry, and biodistribution. Incorporating route of administration as a variable in machine learning frameworks for LNP design may further accelerate the rational development of next-generation therapeutic delivery systems.

We also demonstrated that LNP^cat^ achieved enhanced pancreas expression over LNP^neut^ when delivered either via IV or gastric administration; furthermore, LNP^cat^ outperformed both its neutral analogue in the gastric route (27% increase) and its own intravenous delivery (90% increase), highlighting the synergistic influence of nanoparticle chemistry and route of administration on organ tropism. These results are congruent with findings for intraperitoneal administration^19^. Biodistribution trends from LNP chemistry modifications observed with intravenous delivery were preserved when comparing SM-102 neutral and cationic counterparts delivered intragastrically. Using a Cre recombinase reporter system, we confirmed functional delivery across multiple pancreatic cell types, including acinar and endothelial cells. Targeting of these cell types with LNP^cat^ may enable gastric delivery of therapeutics for a multitude of pancreatic diseases. For example, differentiation of acinar cells to β-like cells that produce insulin can be achieved by inhibiting focal adhesion kinase (FAK)^35^ with siRNAs. It is also possible to achieve β-like cell differentiation from acinar cells by triple transfecting them with MafA, Pdx1, and Ngn3 (3TF) mRNA^36^. Targeting acinar and other cell types in the pancreas may also be useful for other disease settings such as pancreatic cancer. Localized pro-inflammatory cytokines/chemokines could be delivered to a pancreatic tumor. Additionally, strategies to deliver pro-drugs that enable localized cytotoxic effects can be employed, such as delivering mRNAs coding for purine nucleoside phosphorylase (PNP), an Escherichia coli enzyme that converts fludarabine into 2-fluoroadenine, a potent cytotoxic drug^37^.

Together, these findings point toward the development of gastric-interfacing devices capable of delivering lipid nanoparticle payloads for the treatment of pancreatic pathologies, provide a patient preferred administration route that promotes extra-hepatic targeting, and motivate future high-throughput LNP screening efforts to systematically map chemistry–route interactions.

## Methods

### mRNA synthesis and purification

aNLuc and Cre mRNA were prepared as described previously^38,39^. Sequences of GPI-anchored nanoluciferase were ordered as an IDT DNA gBlock. The sequences of the GPI anchor and nanoluciferase have been reported in a previous study^29^. Using a three-molar excess of insert, the gBlock was cloned into a PCR amplified pMA7 vector. The resulting transcripts were purified using a 0.8% agarose gel and verified by Sanger sequencing. In vitro transcription was carried out overnight at 37°C using HiScribe T7 (New England BioLabs) according to the manufacturer’s instructions. The template was then removed by DNase I treatment. To finalize the mRNA preparation, a Cap1 structure was added following denaturation at 65°C for 10 minutes, and the transcripts were enzymatically polyadenylated. The purified RNA products were analyzed by gel electrophoresis to confirm purity.

### Lipid nanoparticle formulation

LNPs carrying aNLuc and Cre mRNA were formulated using the NanoAssemblr Ignite (Precision NanoSystems). SM-102 was purchased from Organix Inc, and cholesterol, PEG lipids and helper lipids were purchased from Avanti Lipids. Aqeuoue (25mM acetate) and organic (100% ethanol) phases were mixed at a flow rate ratio of 3:1 and total flow rate of 12 mL min^-1^.

### Nanoparticle characterization

Nanoparticles were dialyzed using 100 kDa centrifugal filters into 10mM Tris, then sterile purified through 0.22 μm filter. The hydrodynamic diameter of the LNPs was measured using dynamic light scattering (DynaPro Plate Reader III, Wyatt), and the encapsulation efficiency was measured using RiboGreen assay (Invitrogen).

### GI mRNA delivery

All animal procedures were approved by the Georgia Institute of Technology Institutional Animal Care and Use Committee and conducted in accordance with Georgia Institute of Technology animal facility guidelines. C57BL/6J and Ai14 mice were purchased from The Jackson Laboratories. Sprague Dawley rats were purchased from Charles River Laboratories.

Intragastric stomach and colonic Injections: 6-8 week old mice were given one dose of ketoprofen 5 mg kg^-1^ subcutaneously (s.c.) during surgical prep period. If they appeared to be in pain, ketoprofen was continued once a day for the next 2–3 days. Mice were anesthetized with isoflurane in an induction box and maintained on isoflurane for the duration of the procedure via a nose cone (2–3% isoflurane in oxygen for maintenance). The animal was placed in dorsal recumbency, and the abdomen was shaved from just cranial to the xiphoid process to just caudal to the umbilical area. The skin was aseptically prepared with alternating cycles of betadine and 70% ethyl alcohol. The animal was maintained on a heating pad (warm water circulating) and monitored for depth of anesthesia by quality of respiratory effort and response to toe pinch. A 1-cm incision was made on the ventral midline through both the skin and the peritoneal sac. Using atraumatic forceps and sterile cotton tip applicators, we stabilized the area of interest (stomach or colon – distal to the caecum). Using a 30-gauge needle, we injected the lipid nanoparticles suspended in sterile 10 mM Tris buffer into the subserosal side targeting the submucosal space. We then delivered 50-100 µL of fluid based on the lipid nanoparticle concentration. To deliver this amount, we performed up to 10 injections per mouse (forming 10-15 µL boluses) in different locations of the tissue. The abdominal wall was closed with 5-0 PDS or similar absorbable monofilament using a simple interrupted pattern. The skin was closed with a wound clip. Before recovery, mice were given sterile warm 0.9% NaCl s.c. at 20 mL kg^-1^. Mice were given buprenorphine sustained release at 1 mg kg^-1^ s.c. before recovery. Mice were recovered in a cage warmed by a heat lamp and monitored daily.

Buccal and rectal injections: 6-8 week old mice were anesthetized with isoflurane in an induction box and maintained on isoflurane for the duration of the procedure. Using a 30-gauge needle, we injected lipid nanoparticles (50 - 100 µL) submucosally into the buccal tissue underneath the cheek. Additionally, in other groups, we injected lipid nanoparticles by inserting a 30-gauge needle into the rectal lining and injecting 50 – 100 µL slowly. Mice were recovered in a cage warmed by a heat lamp and monitored daily.

Intraduodenal small intestine injections: Sprague Dawley rats (200-300g, Charles River) were given one dose of ketoprofen 5 mg kg^-1^ s.c. during surgical prep period. Rats were anesthetized with isoflurane in an induction box and maintained on isoflurane for the duration of the procedure via a nose cone (2–3% isoflurane in oxygen for maintenance). The animal was placed in dorsal recumbency, and the abdomen was shaved from just cranial to the xiphoid process to just caudal to the umbilical area. The skin was aseptically prepared with alternating cycles of betadine and 70% ethyl alcohol. The animal was maintained on a heating pad (warm water circulating) and monitored for depth of anesthesia by quality of respiratory effort and response to toe pinch. A 1-cm incision was made on the ventral midline through both the skin and the peritoneal sac. Using atraumatic forceps and sterile cotton tip applicators, we stabilized the small intestine immediately distal to the pylorus. We then delivered 50-100 µL of fluid based on the lipid nanoparticle concentration. To deliver this amount, we performed up to 10 injections per mouse (forming 10-15 µL boluses) in different locations of the tissue. The abdominal wall was closed with 5-0 PDS or similar absorbable monofilament using a simple interrupted pattern. The skin was closed with a wound clip. Before recovery, rats were given sterile warm 0.9% NaCl s.c. at 20 mL kg^-1^. Rats were given buprenorphine sustained release at 1 mg kg^-1^ s.c. before recovery. Rats were recovered in a cage warmed by a heat lamp.

### Luciferase assay

16 hours post-injection, mice were sacrificed and 10 organs (pancreas, spleen, kidney, liver, lung, heart, lymph node, buccal, small intestine, colon, stomach, and rectum) were extracted and incubated with 50 mg mL^-1^ Nano-Glo in PBS (Promega) for 20 min, shaking at 800 RPM in a 24-well plate. After incubation, IVIS imaging (IVIS Spectrum CT, PerkinElmer) was performed. Tissues were placed in Nano-Glo Luciferase Assay substrate (Promega) in 24-well plates for 10 min shaking at 100RPM, then measured. Bioluminescence was measured using the IVIS imaging system. Quantitative analysis was performed in Living Image software (PerkinElmer). Images were loaded as a group and brought onto the same scale. The same size ROIs was applied to all organs for analysis. Percentage pancreas delivery in organ screen was determined using the following equation for 10 organs across five routes of administration.

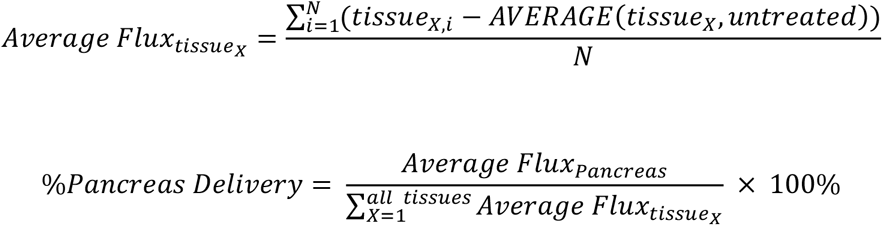

N = number biological replicates

### Tissue sectioning and imaging

Ai14 mice were injected via IV and stomach injections with PBS or 1 mg/kg of Cre mRNA encapsulated in LNP^cat^. Five days later, mice were sacrificed via isoflurane overdose and perfused with PBS and 10% Formalin. Pancreas and liver were extracted and submerged in 10% PFA for 48 h at 4 °C. After that, tissues were washed with 1× PBS 3 times for 15 min at room temperature and embedded in paraffin. Tissues were sliced to 50 μm using VT1000 S Vibratome (Leica). Sections were mounted on Superfrost Plus Microscope Slide (Thermo Fisher Scientific). Sections were generated for IF and H&E stains. To slides stained with H&E, Vestashield Vibrance antifade mounting medium (Vector laboratories) was added, then cover slip added. Imaging was performed using a spinning disk microscope (Nikon) at 10x zoom.

### Serum cytokines and blood chemistry

Mouse sera were sent to IDEXX BioAnalytics and tested on the MILLIPLEX® Mouse Cytokine/Chemokine Magnetic Bead Panel - Premixed 25 Plex (Millipore, cat. no. MCYTMAG70PMX25BK) according to the manufacturer’s protocol. Data were collected by xPONENT 4.3 (Luminex), and data analysis was completed using BELYSA 1.1.0 software. The data collected by the instrument software are expressed as median fluorescence intensity. Standard tox was performed by IDEXX to determine liver enzyme levels.

### Statistics

All statistics were performed in excel or Prism (Graphpad, Version 10). Tests include One-Way ANOVA with Tukey’s correction and Two-Way ANOVA with Tukey’s correction. A minimum of three biological replicates were analyzed for quantification, except PBS groups (minimum of two). All data are presented as the mean ± standard error of mean.

## Acknowledgements

We are grateful to all members of the Dahlman and Abramson Labs for their helpful discussions. We thank Richard Noel and Laura O’Farrell for their help with animal experiment design. We thank the Wallace H. Coulter Department of Biomedical Engineering, Parker H. Petit Institute for Bioengineering and Bioscience, School of Chemistry and Biochemistry, and the Office of the Executive Vice President for Research for their generous support. This work was performed in part at the Georgia Tech Institute for Matter and Systems, a member of the National Nanotechnology Coordinated Infrastructure (NNCI), which is supported by the National Science Foundation (Grant ECCS-2025462).

## Author Contributions

A.M.A., R.G., A.A., and J.E.D created the idea and designed the experiments supporting the investigation of GI-delivered LNPs. A.M.A. and S.R. performed the LNP formulations and characterization. D.S. assisted with laparotomy experiments. A.M.A. and A.S. performed Immunofluorescence. A.A. and J.E.D. provided funding and supervised the project. All authors contributed to writing and editing the manuscript.

## Data Availability

All data associated with this study are present in the paper or the Supplementary Materials.

