## Supplementary for "Gastrointestinal delivery of mRNA lipid nanoparticles selectively targets the pancreas"

**The PDF file includes:**

Figs. S1 to S7

**a**

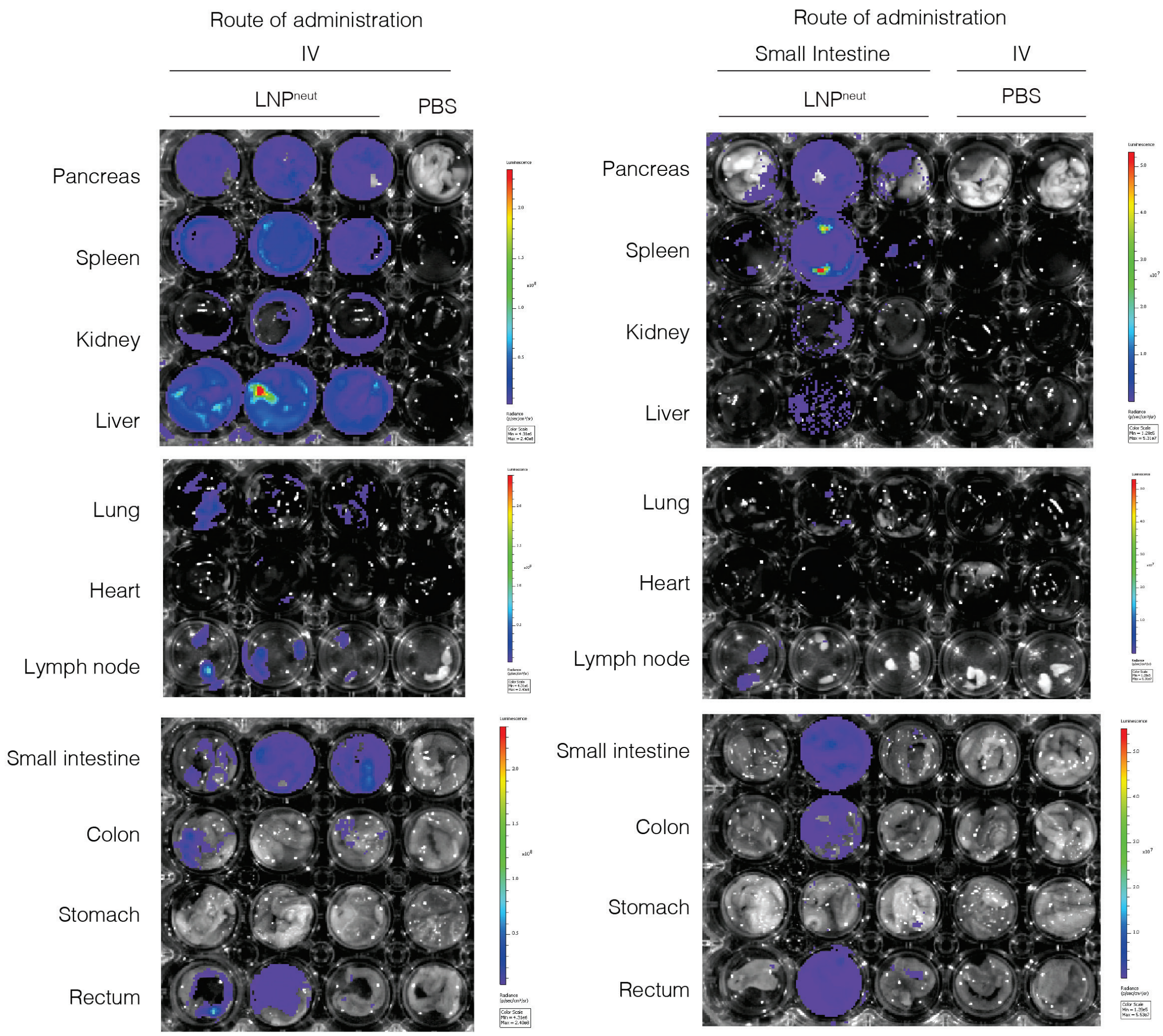

Route of administration
IV
LNP^neut
PBS
Pancreas
Spleen
Kidney
Liver
Lung
Heart
Lymph node
Small intestine
Colon
Stomach
Rectum
Route of administration
Small Intestine
IV
LNP^neut
PBS
Pancreas
Spleen
Kidney
Liver
Lung
Heart
Lymph node
Small intestine
Colon
Stomach
Rectum


**b**

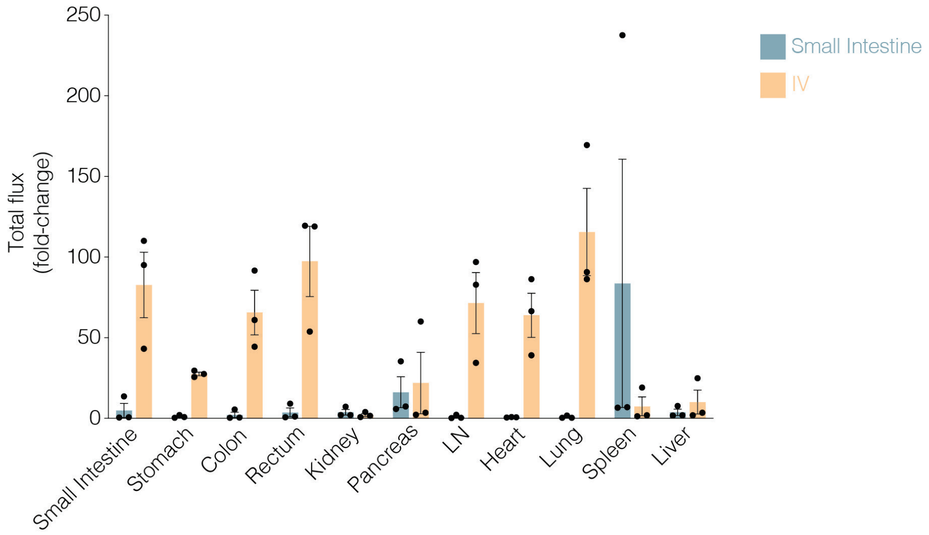

Total flux (fold-change)
250
200
150
100
50
0
Small Intestine
IV
Small Intestine
Stomach
Colon
Rectum
Kidney
Pancreas
LN
Heart
Lung
Spleen
Liver

**Figure S1.** IVIS images after delivering aNLuc mRNA with $LNP^{neut}$ via the small intestine in Sprague Dawley rats. aNLuc RNA was administered at a dose of 0.3 mg $kg^{-1}$ through (a) IV and small intestine. (b) Total flux fold-change. (n = 3 biologically independent animals, Mean ± SEM). PBS-treated rats were used as negative controls and baseline for fold-change calculations.

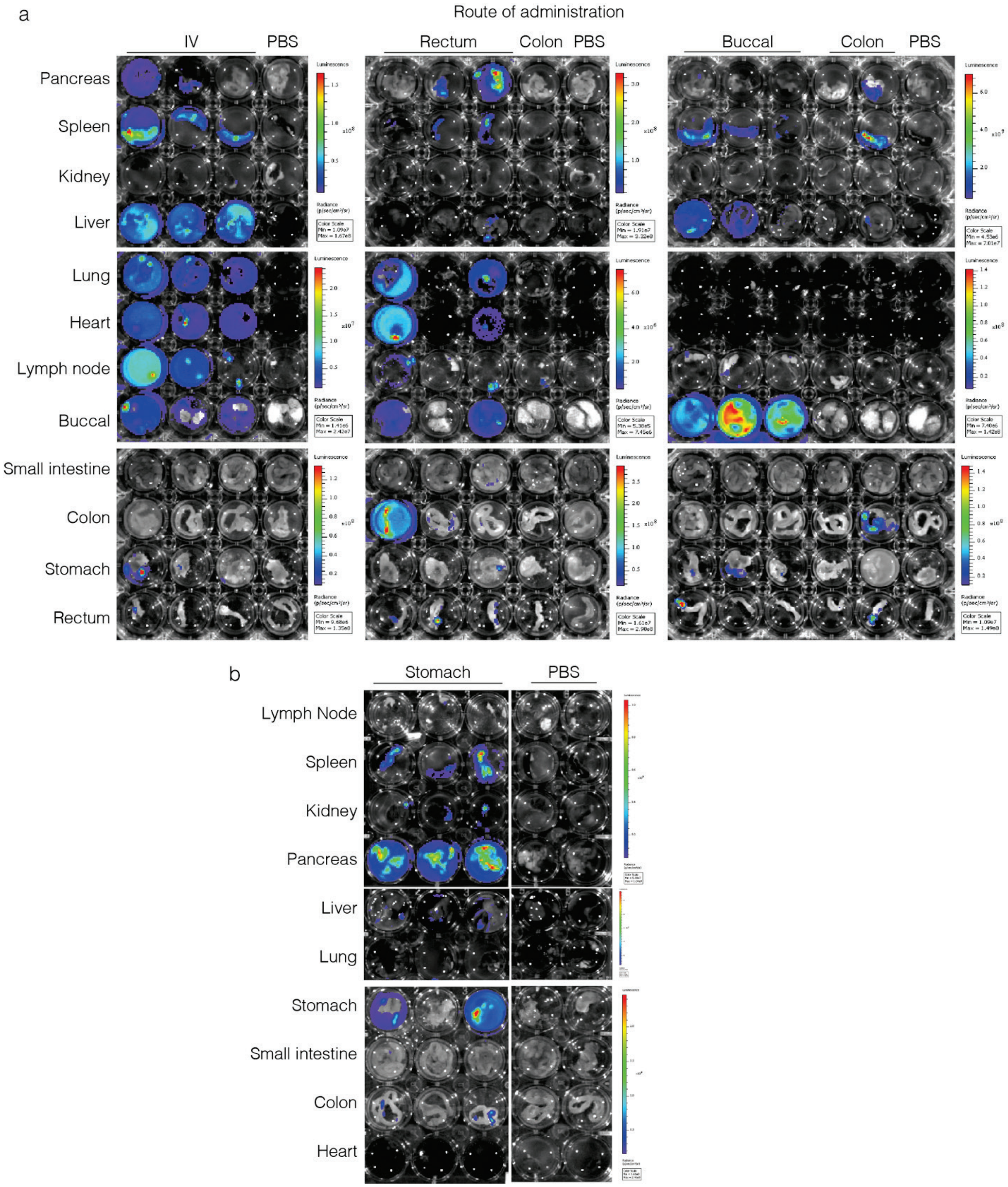


**Figure S2.** IVIS images of GI-organ delivery screen after delivering aNLuc mRNA with LNP$^{neut}$. aNLuc mRNA was administered at a dose of 1 mg kg$^{-1}$ through (a) IV, buccal, colon, rectum, and (b) stomach routes of administration. Luciferase expression was determined via IVIS imaging 18h following dosing. (n = 3 biologically independent animals). PBS-treated mice were used as negative controls and baseline for fold-change calculations.

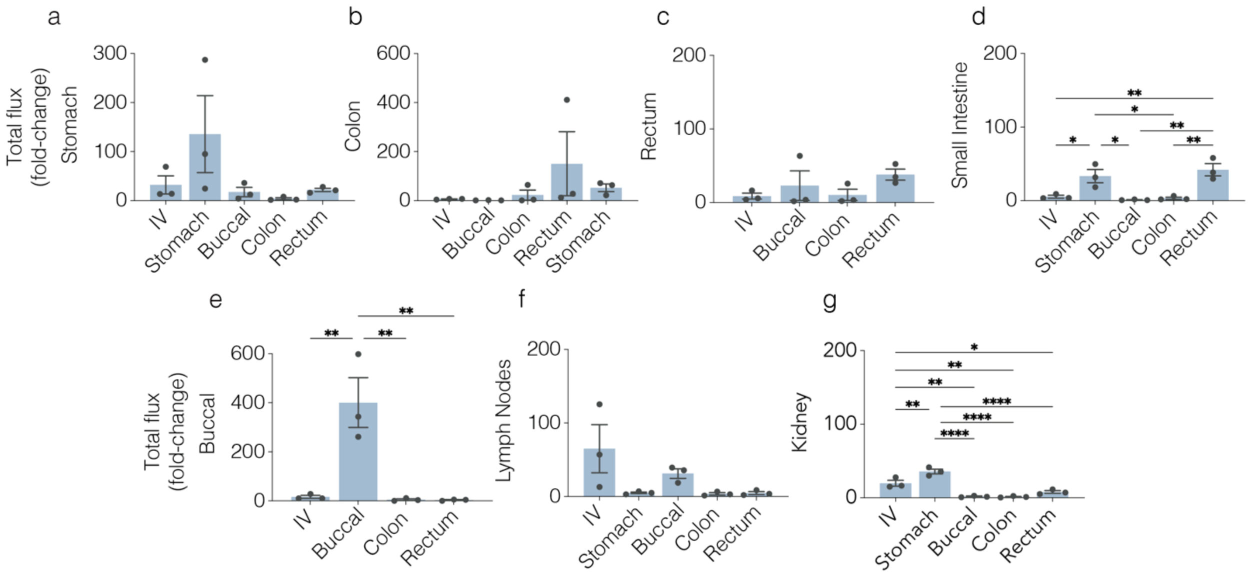


**Figure S3.** Total flux (fold change) for additional organs in GI-organ delivery screen. a) aNLuc RNA was administered at a dose of 1 mg kg$^{-1}$ through multiple routes of administration: intravenously, and to 4 locations across the GI tract - buccal, colon, rectum, and stomach submucosa tissue. Luciferase expression was determined via IVIS imaging at 18h following dosing. (n = 3 biologically independent animals). Quantification of total flux (fold-change) of pancreas across 5 routes of administration compared to PBS-treated group shows stomach with the highest fold-change. (n = 3-4 mice. Individual data points, Mean ± SEM. One-Way ANOVA with Tukey's correction. *p<0.05; **p<0.01; ****p<0.0001.)

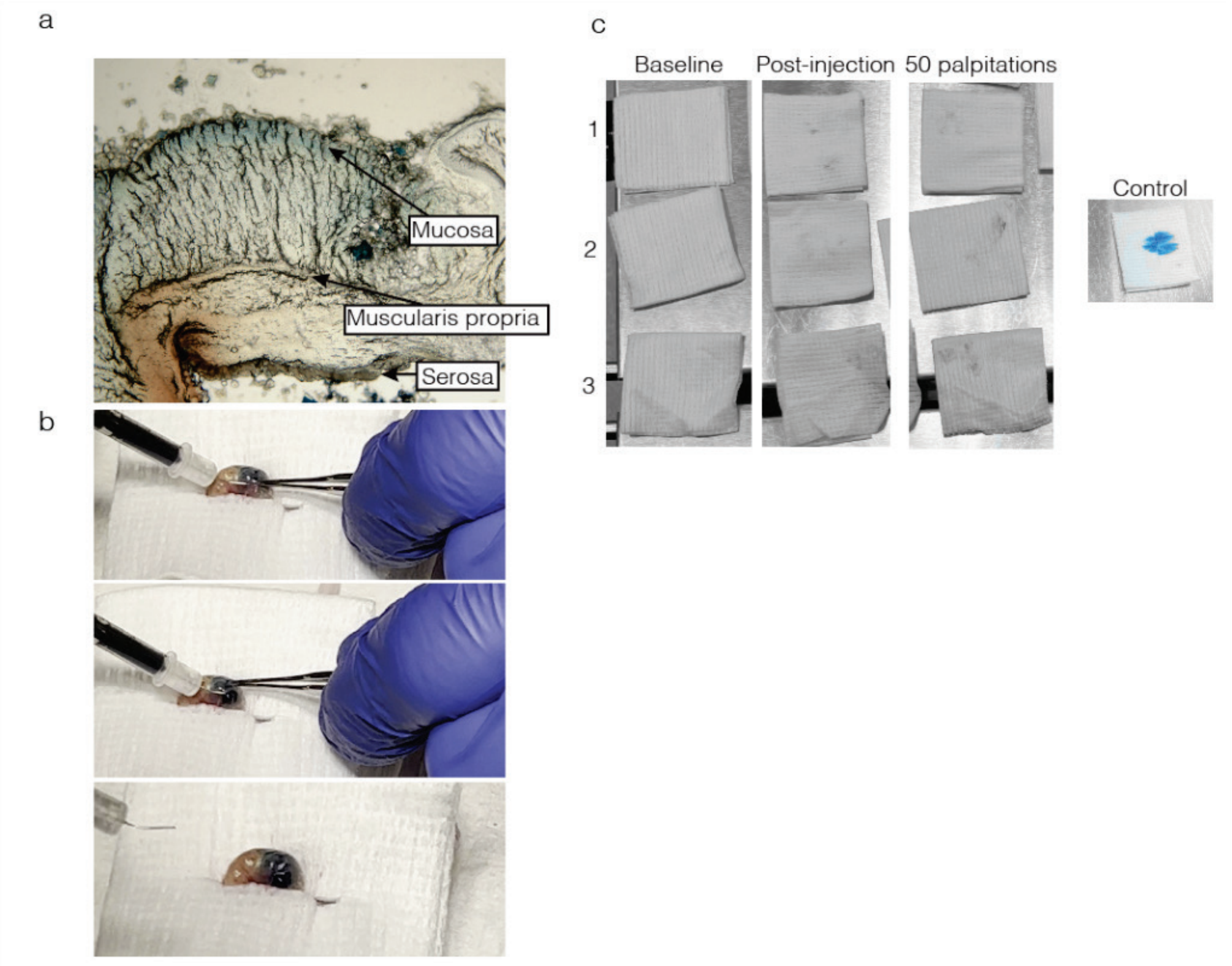


**Figure S4.** Testing GI injection leakage with methylene blue to ensure no leakage. (a) Representative histology of 50 μm thick cross-section of murine stomach after subserosal injections targeting submucosa with 33G needle. (b) Representative images of injection technique. (c) Dry gauze applied to stomach: baseline-before injection, post-injection of 10 μL methylene blue, and after 50 palpitations of abdomen to agitate the affected area. The control gauze was sprayed onto with 10 μL of methylene blue.

a

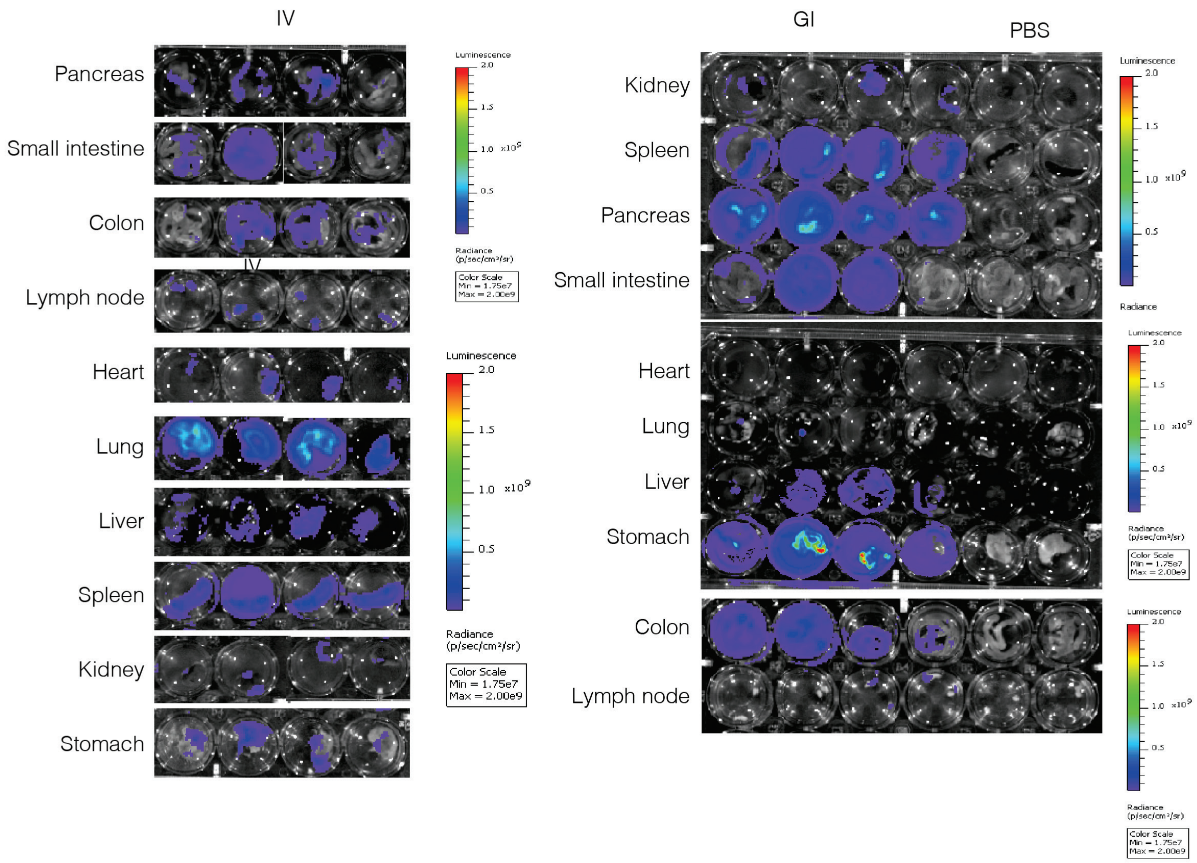


**Figure S5**. Images after administering aNLuc mRNA with LNP$^{cat}$ to stomach and intravenously. aNLuc mRNA encapsulated in LNP$^{cat}$ was delivered via IV or stomach at a dose of 1 mg kg$^{-1}$. Luciferase expression was determined via IVIS imaging 18h following dosing. a-d) images shown for (a) pancreas, (b) liver, (c) heart, and (d) lung. (n = 3 biologically independent animals).

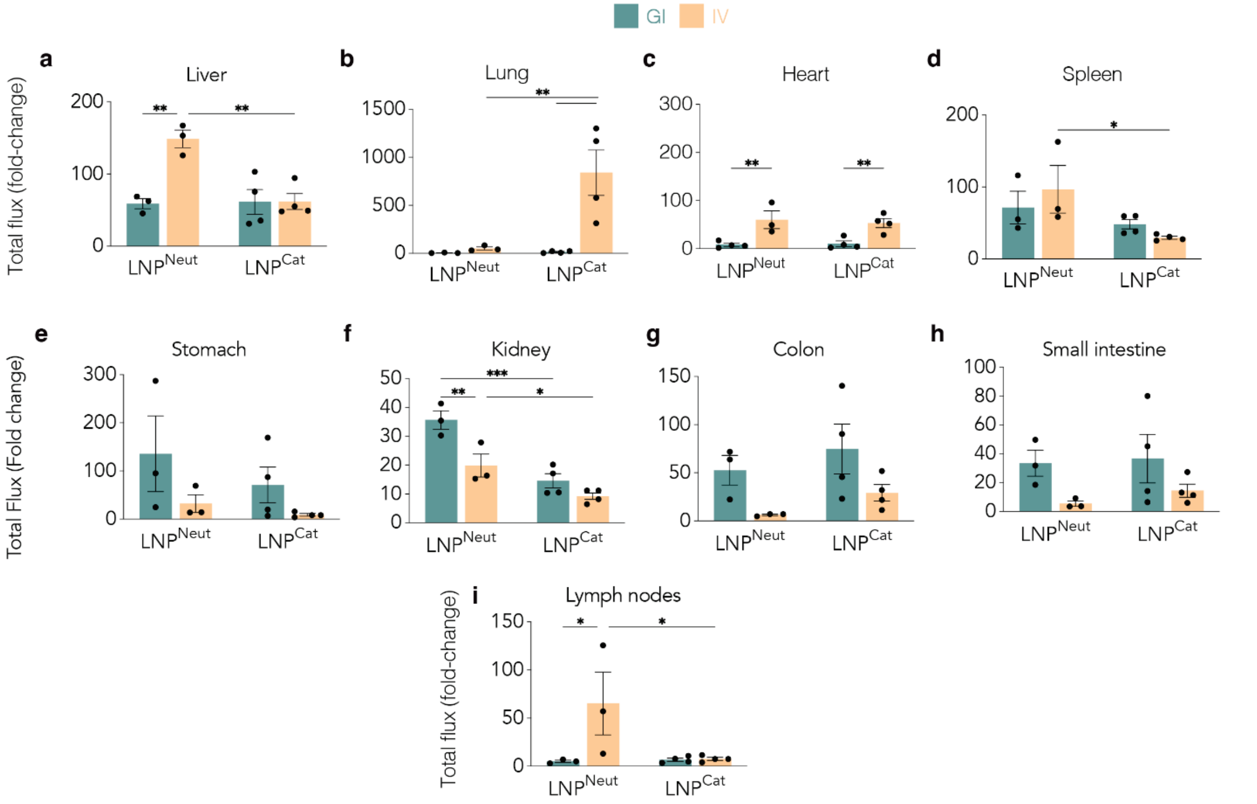


**Figure S6.** Total flux (fold change) for additional organs with aNLuc mRNA with LNP$^{cat}$. a-i) aNLuc mRNA encapsulated in LNP$^{cat}$ was delivered via IV or stomach at a dose of 1 mg kg$^{-1}$. Luciferase expression was determined via IVIS imaging at 18h following dosing. (n = 3 biologically independent animals). Quantification of total flux (fold-change) of (a) liver, (b) lung, (c) heart, (d) spleen, (e) stomach, (f) kidney, (g) colon, (h) small intestine, and (i) lymph nodes across IV and stomach administered mRNA with LNPneut vs LNPcat compared to PBS-treated group. (n = 3-4 mice. Individual data points, Mean ± SEM. Two-Way ANOVA with Tukey's correction. *p<0.05; **p<0.01; ****p<0.0001.)

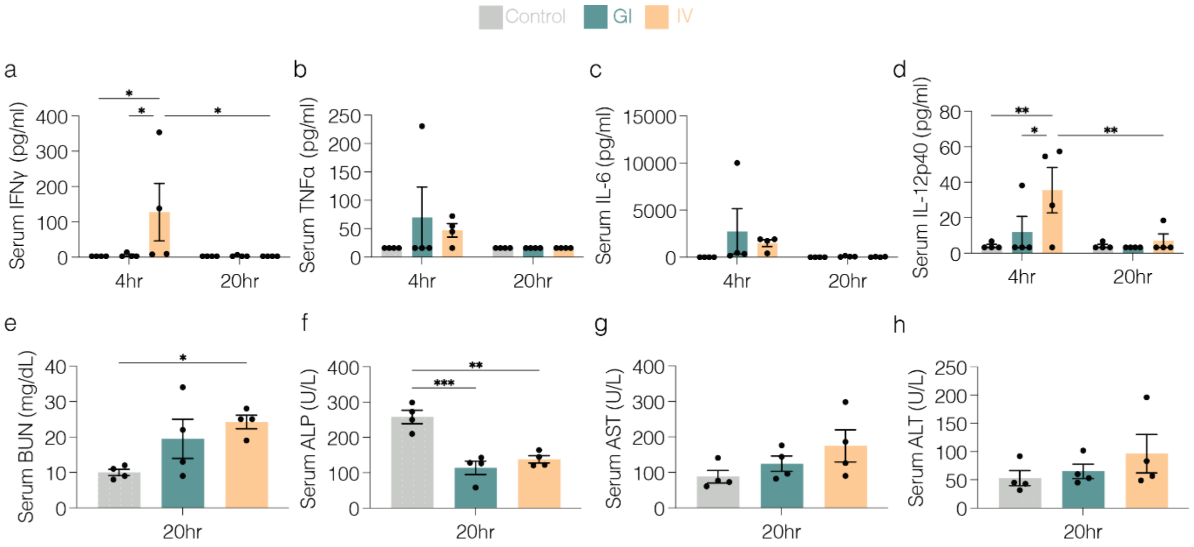


**Figure S7.** Safety profile following GI and IV injections of 3 mg kg$^{-1}$ aNLuc mRNA with LNP$^{cat}$. Mice were injected with 3 mg kg$^{-1}$ mRNA encapsulated in LNP$^{cat}$, and serum was collected 4 hr and 20 hr post injection. a-d) serum cytokines at 4 hr and 20 hr quantifying (a) IFNγ, (b) TNFα, (c) IL-6, and (d) IL-12p40. e-h) Blood chemistry was performed on serum to quantify liver enzymes at 20hr: (e) BUN, (f) ALP, (g) AST, (h) ALT. (n = 3-4 mice. Individual data points, Mean ± SEM. One-Way ANOVA with Tukey's correction. *p<0.05; **p<0.01; ****p<0.0001.)
